# Expression of novel androgen receptors in three GnRH neuron subtypes in the cichlid brain

**DOI:** 10.1101/2024.02.02.578641

**Authors:** Mélanie Dussenne, Beau A. Alward

## Abstract

Within a social hierarchy, an individuals’ social status determines its physiology and behavior. In *A. burtoni*, subordinate males can rise in rank to become dominant, which is accompanied by the upregulation of the entire HPG axis, including activation of GnRH1 neurons, a rise in circulating androgen levels and the display of specific aggressive and reproductive behaviors. Cichlids possess two other GnRH subtypes, GnRH2 and GnRH3, the latter being implicated in the display of male specific behaviors. Interestingly, some studies showed that these GnRH neurons are responsive to fluctuations in circulating androgen levels, suggesting a link between GnRH neurons and androgen receptors (ARs). Due to a teleost-specific whole genome duplication, *A. burtoni* possess two AR paralogs (ARα and ARβ) that are encoded by two different genes, *ar1* and *ar2*, respectively. Even though social status has been strongly linked to androgens, whether ARα and/or ARβ are present in GnRH neurons remains unclear. Here, we used immunohistochemistry and *in situ* hybridization chain reaction (HCR) to investigate *ar1* and *ar2* expression specifically in GnRH neurons. We find that all GnRH1 neurons intensely express *ar1* but only a few of them express *ar2*, suggesting the presence of genetically-distinct GnRH1 subtypes. Very few *ar1* and *ar2* transcripts were found in GnRH2 neurons. GnRH3 neurons were found to express both *ar* genes. The presence of distinct *ar* genes within GnRH neuron subtypes, most clearly observed for GnRH1 neurons, suggests differential control of these neurons by androgenic signaling. These findings provide valuable insight for future studies aimed at disentangling the androgenic control of GnRH neuron plasticity and reproductive plasticity across teleosts.

## 1 Introduction

Social species often organize into dominance hierarchies in which social rank greatly influences individuals’ physiological and behavioral traits ^1^. In these hierarchies, dominant (DOM) and subordinate (SUB) individuals coexist and greatly differ in their physiology and behaviors. DOM animals monopolize territory and food resources, mate frequently, and maintain their dominance status through intimidation and aggression. On the contrary, SUB individuals perform submissive behaviors, have limited access to resources, and mate very infrequently if at all.

In some species, DOM and SUB individuals can switch status depending on social context. This is the case in the African cichlid fish *Astatotilapia burtoni*, in which a male-based social hierarchy includes two distinct socially-mediated phenotypes ^2–7^. DOM males typically possess large testes and high circulating sex steroids levels, promoted by the up-regulation of the Hypothalamo-Pituitary-Gonadal (HPG) axis ^8,9^. On the contrary, SUB males, which constantly monitor the social environment awaiting an opportunity to rise in rank ^10–12^, present a suppressed HPG axis. As soon as SUB males ascend socially, their entire HPG axis is activated. Hypophysiotropic gonadotropin-releasing hormone (GnRH1) neurons, located in the preoptic area, stand at the apex of the HPG axis ^13^ and show incredible plasticity during social ascent in male *A. burtoni*. For example, after perception of social opportunity GnRH1 mRNA levels increase in minutes and GnRH1 neurons progressively increase in size to reach that of DOM males within just a few days ^14^. GnRH1 neurons in DOM males have distinct electrical properties compared to those in SUB males ^15^. These physiological changes in GnRH1 neurons reverberate along the HPG axis, increasing levels of circulating sex steroid levels as well as testes mass in approximately two weeks following social opportunity ^9^.

In teleosts, other forms of GnRH exist. For instance, in cichlids, distinct genes encoding both GnRH2 and GnRH3 are present as well. GnRH2 and GnRH3 neurons are located in the midbrain and olfactory bulbs, respectively ^16–20^. In contrast to GnRH1 neurons, GnRH2 and GnRH3 neurons do not appear to regulate reproductive physiology. In teleosts, GnRH3 neurons have been linked to the expression of mating behaviors in males ^21,22^ and females ^23^. The function of GnRH2 neurons is still unclear but available evidence suggest they may regulate energy balance ^24^.

Androgens have been shown to affect the plasticity of GnRH neurons in teleost fishes. For example, in male *A. burtoni*, castration, which leads to the dramatic reduction of circulating androgens, leads to GnRH1 neuron hypertrophy, a change that is reversed upon androgen treatment ^25,26^. In adult female Mozambique tilapia (*Oreochromis mossambicus*), treatment with the androgens 11-KT or methyltestosterone, but not 17β-estradiol (E2), resulted in enhanced GnRH3 neurogenesis and was associated with an increase in male-typical nest building behavior ^27,28^. 11-KT is a non-aromatizable androgen, indicating that the effects of androgens on GnRH1 and GnRH3 neurons are likely due to actions at androgen receptors (ARs), either directly in GnRH neurons or through an indirect mechanism. Complicating this question further is the presence of novel AR paralogs encoded by distinct genes in most teleost fishes due to a teleost-specific whole-genome duplication ^29,30^. Indeed, while all other vertebrates have a single *ar* gene encoding one AR, most teleost fishes have two distinct *ar* genes—*ar1* and *ar2—*encoding two different ARs, ARα and ARβ, respectively ^31–33^.

The presence of androgen receptors (ARs) in GnRH neurons in teleost fishes has not been definitively established. In *A. burtoni*, there is some evidence that both *ar* genes may be present in GnRH1 neurons but the work supporting this did not confirm this beyond a single plane of imaging—that is, z-stacks or a similar method were not used to confirm the presence of signal for both genes in the same plane and thus in the same cell ^34^. In a recent study in Nile tilapia (*Oreochromis niloticus*) results of qPCR for *ar* genes in dissociated GnRH neurons suggest that only *ar2* is expressed in all GnRH subtypes ^19^. However, these results were not confirmed spatially in intact tissue. Therefore, it is not definitively known whether ARs are present in GnRH neurons in the teleost brain.

Here, we used *in situ* hybridization chain reaction (HCR), immunohistochemistry, and a novel approach combining these techniques to characterize *ar1* and *ar2* expression in GnRH neurons. We discovered that while both *ar1* and *ar2* are expressed in GnRH neurons, differences between the expression of *ar* paralogs in all three GnRH neuron subtypes exist. Our findings have important implications for disentangling the extraordinary reproductive plasticity among teleost fishes.

## 2 Material and Methods

### 2.1 Ethics

Experimental procedures were conducted according to the ethical guidelines for the care and use of laboratory animals. All experiments were done in accordance with The University of Houston Institutional Animal Care and Use Committee (Protocol #202000001).

### 2.2 Animals

Individuals used in these experiments were adult *A. burtoni* derived from a wild-caught population from Lake Tanganyika, Africa ^35^. They were maintained in different recirculating systems all presenting the same physico-chemical parameters that mimicked their natural equatorial environment (28°C, pH = 8.2, photoperiod 12 h:12 h (L:D), under constant aeration). Aquaria contained gravel-covered bottoms with terracotta pots cut in half to serve as shelters and spawning territories.

Fish were fed *ad libitum* during the day with commercial cichlid pellets (Ken’s Premium Cichlid Pellets 1.5, KensFish, Taunton, MA, USA) and supplemented with brine shrimp (Bio-Pure Frozen Brine Shrimp, Hikari Sales USA, Inc.).

Fry were collected between 0 and 12 days post fertilization and placed in an shaking incubator until their yolk sac was fully resorbed. Then, larvae were transferred into tanks in one of the recirculating systems, where they were fed with powdered pellets supplemented with Freeze-dried cyclops (San Francisco Bay Brand, Inc., Newark, CA) until they were large enough to feed on pellets.

In this study, WT fish were used, as well as GnRH1:eGFP transgenic fish and ARα and ARβ knockouts for validation purposes. Methods used to generate GnRH1:eGFP transgenic as well as ARα and ARβ knockouts have already been thoroughly described (for more details, see ^36,37^).

To determine genotype, a small piece of the caudal fin of juvenile fish was collected for DNA isolation and PCR. Briefly, a small portion (∼1-2mm) of the caudal fin was excised and placed in a PCR tube. DNA was extracted by adding 180 μl of NaOH (50 mM) in each tube which was then incubated at 94°C for 15 min. Then, 20 μl of Tris-HCl (pH 8) was added to each sample which was then vortexed, centrifuged and incubated at 4°C for 1 h. Then, amplicons spanning the AR mutation target or the GnRH1:eGFP transgene sites were PCR-amplified using their specific primers (see Supplementary Table 1). PCR products were visualized on a gel. To check for the ARβ mutation, samples were Sanger-sequenced using the ARβFlankF primer (Lone Star Labs, Houston, TX). Fish were then housed with individuals of the same genotype.

### 2.3 Dissection and tissue collection

Fish were euthanized by immersion in an ice water bath, subsequently weighted (body mass, BM) and measured. After cervical transection, brains were exposed from the olfactory bulbs to the spinal cord and fixed in the head in 4% PFA (pH 7.0 for HCR, pH 7.4 for IHC) for 24 h at 4°C. Heads were then rinsed in 1x PBS (3 × 5 min, then ∼6 h at 4°C) and cryoprotected in 30% sucrose overnight at 4°C. Brains were extracted from the heads, embedded in Neg-50 (22-046-511, Fisher Scientific) and kept at -80°C until sectioning. Gonads were removed from the visceral cavity and weighted (gonads mass, GM) in order to calculate the gonadosomatic index [GSI; (GM/BM)*100].

Brains were sectioned in the coronal plane using a cryostat (Thermo Scientific, HM 525 NX) at either a thickness of 20 μm on three alternate slide series for validation experiments or a thickness of 30 μm on two alternate slide series for experimental individuals. Slides were air-dried for 24 h then stored at -80°C until further processing.

### 2.4 Antibody validations

#### 2.4.1 Anti-LHRH antibody validations

The anti-LHRH antibody (Immunostar) has previously been shown to recognize the three forms of GnRH in Nile tilapia (*Oreochromis niloticus*), a species closely related to *A. burtoni* ^38^ To further assess its specificity in *A. burtoni*, we performed several validation steps. First, to confirm that the anti-LHRH antibody specifically labels GnRH1 neurons, we pre-incubated this primary antibody with the GnRH1 peptide (L7134, Sigma-Aldrich) at increasing concentrations (1, 10, 100 μg/mL). For this purpose, the brain of a GnRH1:eGFP transgenic female was cut into three alternate series. A double IHC was performed (see protocol section 2.5), using 1) the monoclonal mouse anti-eGFP antibody (MA1-952, thermoFisher) and 2) the polyclonal rabbit anti-LHRH antibody pre-incubated with the GnRH1 peptide (one GnRH1 peptide concentration per slide series). To detect these primary antibodies, the Alexa Fluor 488 goat anti-mouse (A-11029, ThermoFisher) and the AF 647 goat anti-rabbit (111-605-003, Jackson ImmunoResearch) secondary antibodies, respectively, were used.

Similarly, to confirm the anti-LHRH antibody also recognizes GnRH2 and GnRH3, we pre-incubated it with the GnRH2 and GnRH3 peptides (SCP0183 and L4897 respectively, Sigma-Aldrich) at a concentration of 100 μg/mL. One female brain was also sectioned into three slide series; one of them was incubated with the anti-LHRH antibody to serve as a positive control, and the two others were incubated with the anti-LHRH antibody pre-incubated with either the GnRH2 or GnRH3 peptides. Additionally, to further confirm the specificity of the LHRH antibody, we performed a simultaneous HCR and IHC using *gnrh1, gnrh2, gnrh3* HCR probes altogether with the LHRH antibody (see protocol section 2.7).

#### 2.4.2 Anti-ARα and anti-ARβ antibodies validations

Two custom polyclonal antibodies that were directed against ARα and ARβ were produced (AvesLabs). The anti-ARα antibody was raised against an ARα synthetic peptide (CFDKLRTSYIKELDRLA SHHGETTRTQR) and the anti-ARβ antibody was raised against an ARβ synthetic peptide (CZQKTEESRVYFTKTPTGSS).

In an attempt to validate the specificity of these antibodies, we first wanted to demonstrate the absence of ARα and ARβ immunolabelling in the brains of ARα and ARβ KO individuals, respectively. For this purpose, we performed a double IHC on the brain of one ARα-KO, one ARβ-KO and one WT individuals, the latter being used as an immunostaining positive control. Antibodies used were 1) the anti-LHRH primary antibody visualized with the Alexa-Fluor 488 secondary antibody and either 2) the anti-ARα or anti-ARβ primaries visualized with the Alexa-Fluor 594 secondary (see IHC protocol section 2.5). To further test for the specificity of these antibodies, we performed a western blot (see Supplementals 1). Several tissue types (ovaries, testes, brain, liver) were used to see whether the custom antibodies could selectively bind to their target when presented to a heterogenous sample.

### 2.5 Immunohistochemistry

Two days before the start of IHC, slides were dried using a desiccator. On the first day of IHC, the antigen retrieval step was performed by immersing the slides in boiling citrate buffer (pH 6, 2 × 5 min). After three rinses in PBS (5 min), slides were blocked with 0.4% Triton in PBS containing 10% normal goat serum (NGS) and 1% milk powder (1 h, RT). Slides were then incubated with the anti-eGFP primary antibody (MA1-952, thermoFisher) at a final dilution of 1/500 in 0.4% PBST containing 0.5% of NGS in a sealed humid chamber (overnight, 4°C).

On day 2, after three rinses in PBS (5 min), slides were incubated with the anti-LHRH primary antibody (20075, lot XX, Immunostar) at a dilution of 1/500 in 0.4% PBST containing 0.5% NGS (overnight, 4°C). On day 3, after three rinses in PBS (5 min), slides were incubated with both secondary antibodies AF 488 goat anti-mouse (A-11029, ThermoFisher) and AF 647 goat anti-rabbit (111-605-003, Jackson ImmunoResearch) at a dilution of 1/300 in 0.4% PBST containing 0.5% NGS (1 h, RT in the dark). Slides were finally rinsed in PBS (3x 5 min) and coverslipped using ProLong Gold antifade reagent with DAPI (8961S, Cell Signalling Technology). For the custom antibodies validation, we used the anti-LHRH antibody and either custom anti-AR antibodies (AvesLabs), visualized with the AF 488 goat anti-rabbit and AF 594 goat anti-chicken secondaries (111-545-144 and 103-585-155 respectively, Jackson ImmunoResearch).

### 2.6 *In situ* hybridization chain reaction (HCR)

Two individuals were used for HCR; one WT male and one WT gravid female (GSI= 0.5 and 8.1, respectively). For each individual, the two slides series were double-labelled using 1) the *gnrh1, gnrh2*, and *gnrh3* probes altogether and 2) either the *ar1* or the *ar2* probes.

Slides were removed from the -80°C freezer and exposed to UV light for 1 h at room temperature, then washed in 0.2% PBST (30 min, RT) and fixed in 4% PFA (pH 7.4, 15 min, RT). To remove the Neg-50 from the slides, they were washed through a series of ethanol baths of increasing concentration (50%, 70%, 100%, 5 min each, RT) after which slides were air dried for 5 min. Slides were then incubated in 10 μg/mL proteinase K solution (10 min, 37°C) after which they were immediately rinsed in PBS (2 × 5 min, RT). Slides were pre-hybridized by placing warm probe hybridization buffer onto the slides (Molecular Instruments, 10 min, 37°C). The probe hybridization solution was prepared, with each probe at a final concentration of 16 nM in pre-heated probe hybridization buffer (37°C). The HCR probes used here were 1) *gnrh1, gnrh2, gnrh3* probes all coupled with B3 amplifier and 2) either the *ar1* or *ar2* probe (coupled with B1 and B2 amplifiers, respectively, Molecular Instruments). At the end of the pre-hybridization step, excess buffer was removed from each slide and the probe hybridization solution was added to each slide and covered with a plastic coverslip. Slides were finally placed in a humid sealed chamber at 37°C overnight.

On day 2, plastic coverslips were removed from the slides by immersing them in warm (37°C) probe wash buffer (Molecular Instruments). Sections were then sequentially rinsed in a series of pre-heated (37°C) probe wash solutions of decreasing concentrations (75%, 50%, 25%, diluted in 5x SSCT; 15 min each, 37°C). Slides were rinsed twice in 5x SSCT (15 min at 37°C and 5 min at RT). We then proceeded with the pre-amplification step, for which slides were covered with amplification buffer (Molecular Instruments) and placed in a sealed humid chamber (30 min, RT). During this step, appropriate hairpins (B3-488 and B1-647 or B2-647) were each aliquoted in PCR tubes and activated by being incubated at 95°C for 90 sec using a thermocycler. Hairpins were then allowed to cool down in the dark (30 min, RT). Once the pre-amplification step was completed, the hairpin amplification solution was prepared by adding each activated hairpin (at a final concentration of 60 nM) in amplification buffer. Each slide was covered with this hairpin amplification solution, topped with a plastic coverslip and placed in a sealed humid chamber in the dark overnight at RT.

On day 3, slides were discharged from their plastic coverslip as previously, then rinsed twice in 5x SSCT (2 × 30 min, RT) and twice in PBS (5 min, RT in the dark). We then proceeded to the mounting of the slides with ProLong Gold antifade reagent (9071, Cell Signalling Technology).

### 2.7 Simultaneous HCR and IHC

This protocol was adapted from a recently published protocol 39. This technique was used 1) in order to confirm the specificity of the *gnrh* probes and the anti-LHRH antibody; that is, to confirm that the *gnrh* probes and the antibody effectively labelled the same cells and 2) to simultaneously label GnRH neurons and *ar* mRNA.

The first day of the protocol was similar to the HCR protocol, except that the proteinase K treatment was not applied to the sections. The HCR probes used were the *gnrh1, gnrh2* and *gnrh3* coupled with a B3 amplifier (Molecular Instruments) to detect all three simultaneously.

On day 2, the anti-LHRH antibody (Immunostar) was added to the hairpin amplification solution at the same concentration used in IHC (1/500). As previously, the hairpins used for the *gnrh* probes were B3-488. On day 3, the plastic coverslip was taken off using 5x SSCT, then slides were rinsed in 5x SSCT (3 × 5 min, at RT in the dark) and in PBS (2 × 5 min, RT in the dark). Slides were then incubated with the secondary antibody (AF 647 goat anti-rabbit, 111-605-003, Jackson ImmunoResearch) at a dilution of 1/300 in 0.2% PBST (1 h, RT in a sealed dark chamber). Slides were finally rinsed with PBS (3 × 5 min, RT in the dark) and coverslipped using ProLong Gold antifade reagent with DAPI (8961S, Cell Signalling Technology).

To simultaneously label GnRH neurons and ARs, we used both *ar* probes (coupled with B1-647 and B2-488 hairpins) and the anti-LHRH primary antibody visualized with the AF 405 goat anti-rabbit secondary (111-475-003, Jackson ImmunoResearch).

To image the stained sections, we used a Nikon Eclipse T*i*2 together with the NIS-Elements software (version 5.30.05). Pictures showing the same sections imaged with two different fluorophores always belong to the same z-stack plan (0.3 μm). Validation photomicrographs have been taken with the same exposure settings. All of the findings supporting the claims made in this manuscript are available on the figures in the main text and present in the Supplementary Materials file.

## 3 Results

### 3.1 Antibodies validations

#### 3.1.1 Anti-LHRH antibody

To confirm the anti-LHRH antibody selectively labelled GnRH1 neurons, we immunostained the brain of a GnRH1:eGFP transgenic fish with the anti-eGFP antibody and the anti-LHRH antibody preincubated with different GnRH1 peptide concentrations. Preincubation resulted in the disappearance of GnRH1 immunoreactivity in a concentration-dependent manner (Fig 1A). Immunostaining revealed by the anti-LHRH antibody showed a clear 1:1 correspondence to the results revealed by the anti-eGFP antibody, indicating that the labeled neurons were indeed GnRH1 neurons (Fig 1A).

**Fig 1:**
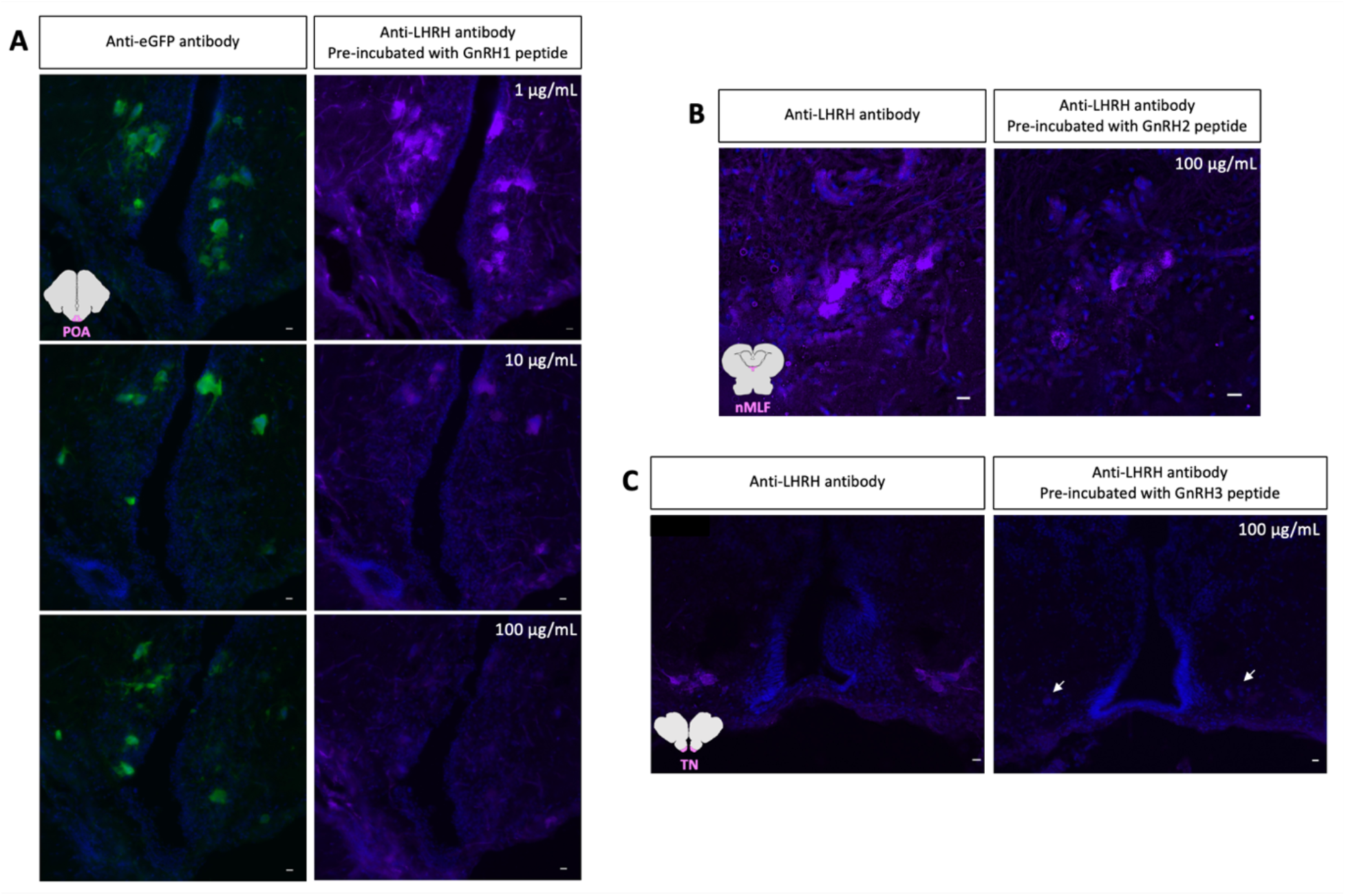
The anti-LHRH antibody specifically recognizes all GnRH variants. A, left column: GnRH1 immunoreactive (-ir) cell bodies stained for eGFP. Right column: same sections showing GnRH1-ir neurons stained using the anti-LHRH antibody pre-incubated with different GnRH1 peptide concentrations. B, GnRH2 neurons on two adjacent sections, immunolabeled with the anti-LHRH antibody (left, control) and with the anti-LHRH antibody preincubated with the GnRH2 peptide (right). C, GnRH3 neurons on two adjacent sections, immunolabeled with the anti-LHRH antibody (left, control) and with the anti-LHRH antibody preincubated with the GnRH3 peptide (right). Blue, DAPI. Scale bars, 10 μm.

Similarly, to confirm the anti-LHRH antibody labelled GnRH2 and GnRH3 neurons as well, we immunostained a WT brain with either the anti-LHRH antibody or the anti-LHRH antibody pre-incubated with either GnRH2 or GnRH3 peptide. Pre-incubation of the antibody with the GnRH2 peptide decreased the immunostaining intensity compared to the control condition (Fig 1B). Pre-incubation of the antibody with the GnRH3 peptide resulted in the total disappearance of immunoreactivity in GnRH3 neurons (Fig 1C).

To further confirm the anti-LHRH antibody immunostains the three forms of GnRH, we performed a simultaneous HCR and IHC, using all three *gnrh* HCR probes together with the anti-LHRH antibody. *gnrh* probes together with the anti-LHRH antibody effectively labelled the same cells, namely the GnRH1, GnRH2 and GnRH3 neuronal populations (Fig 2). Based on previous work 38 together with the present results, we can attest that the anti-LHRH antibody effectively labels the three forms of GnRH in our cichlid fish species.

**Fig 2:**
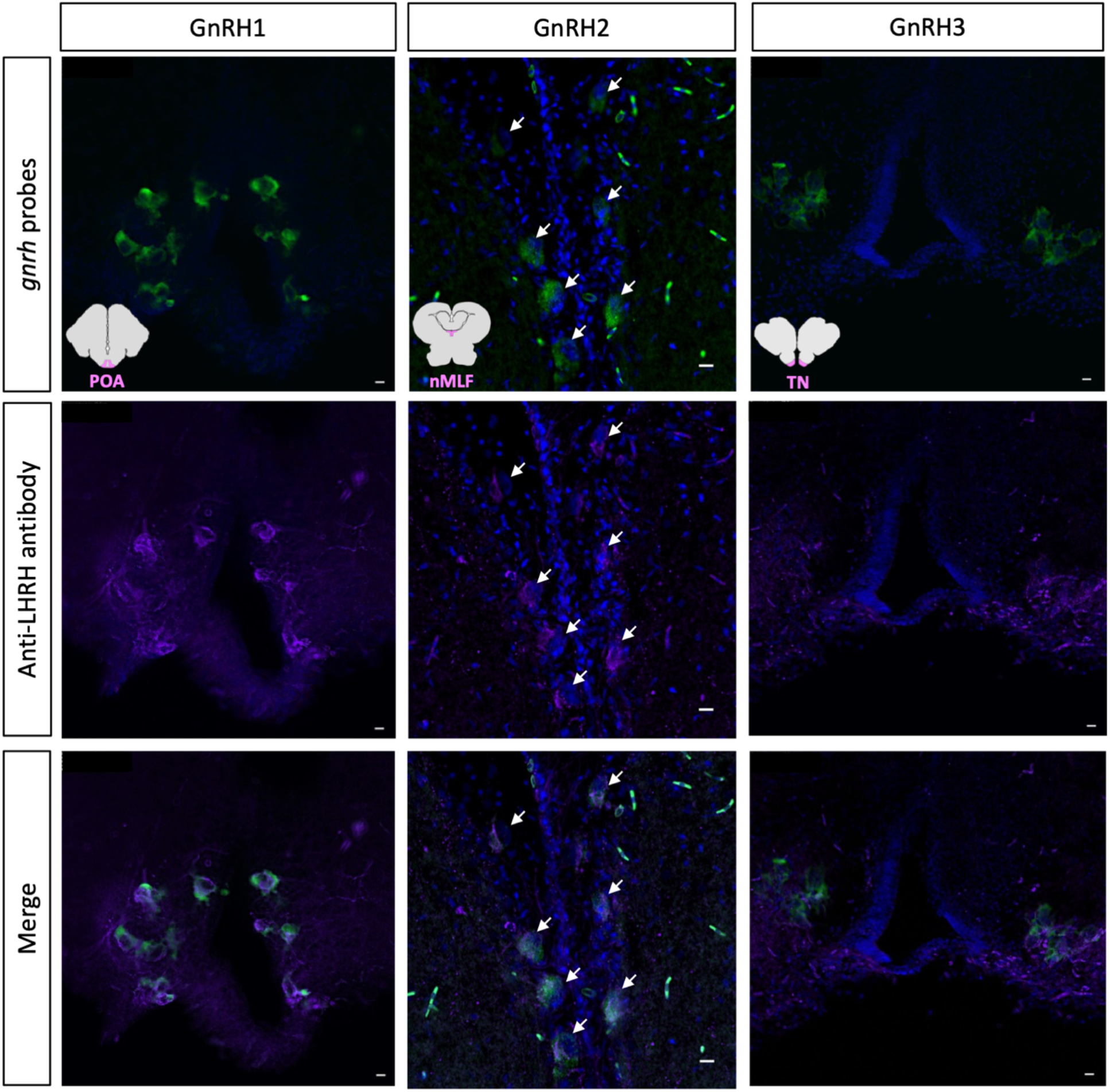
GnRH HCR probes together with the anti-LHRH antibody recognize the same cells. Photomicrographs showing staining of *gnrh* probes used altogether (top row), immunostaining with the anti-LHRH antibody (middle row) and merge of these stainings (bottom row) in GnRH1, GnRH2 and GnRH3 neurons (left, middle and right columns respectively). Blue, DAPI. Scale bars, 10 μm.

#### 3.1.2 Anti-ARα and anti-ARβ custom antibodies

As a first step in our validation process for these antibodies, we used ARα KO and ARβ KO individuals to check for the specificity of the anti-ARα and anti-ARβ antibodies, respectively. As KO individuals do not express the target protein, a specific antibody directed against this protein should not yield any staining. Thus, a WT individual was used as a positive control while the two KO individuals were used as negative controls for their respective antibodies. Unfortunately, the anti-ARα antibody revealed intense immunostaining in the POA (especially in GnRH1 neurons) in the ARα KO individual. Similarly, ARβ immunostaining could be seen in the POA of the ARβ KO individual. Interestingly, the intensity of the immunostainings yielded by theses antibodies in KO individuals was similar to the signaling intensity in the control WT brain (not shown), suggesting these antibodies are not specific.

Therefore, we performed a western blot in order to identify any off-target binding for these antibodies. For this purpose, AR rich tissues (gonads, brain, liver) of several WT, ARα KO and ARβ KO animals were used (see protocol in Supplemental 1). For both anti-ARα and anti-ARβ antibodies, multiple bands were identified for all samples (including KO individuals). Based on these results, we concluded that these antibodies show cross-reactivity with several off-target proteins and that they show poor specificity. As these custom antibodies could not be used any further, we investigated the presence of ARs in GnRH neurons using only *ar* HCR probes.

### 3.2 *ar1* and *ar2* gene expression pattern in GnRH neurons

To investigate whether *ar1* and *ar2* genes are expressed in GnRH neurons, we sampled the brain of two WT animals (1 male, 1 female). For both individuals, brains were sectioned on alternate slide series; one was labelled with *gnrh* and *ar1* HCR probes, the other slide series with *gnrh* and *ar2* HCR probes. Previous work showed that in *A. burtoni*, the preoptic area is rich in *ar1* and *ar2* mRNA 34. This is confirmed in the present study, with both *ar1* and *ar2* genes being intensely expressed in the nPPa, especially along the third ventricle. In our species, GnRH1 neurons were localized in both this *ar1* and *ar2* mRNA rich area (along the third ventricle, see area delimited by the dashed line in Fig 3) and outside of it. Interestingly, in these GnRH1 neurons *ar1* and *ar2* genes show a distinct expression pattern. Specifically, *ar1* gene was expressed in every single GnRH1 neuron, whether they were present in the AR-rich area or not (see white arrows in Fig 3, A and B), while *ar2* mRNA was present in rare GnRH1 neurons inside this AR-rich area (see white arrow in Fig 3 D) but not in GnRH1 neurons outside of this area (see red arrows in Fig 3 D).

**Fig 3:**
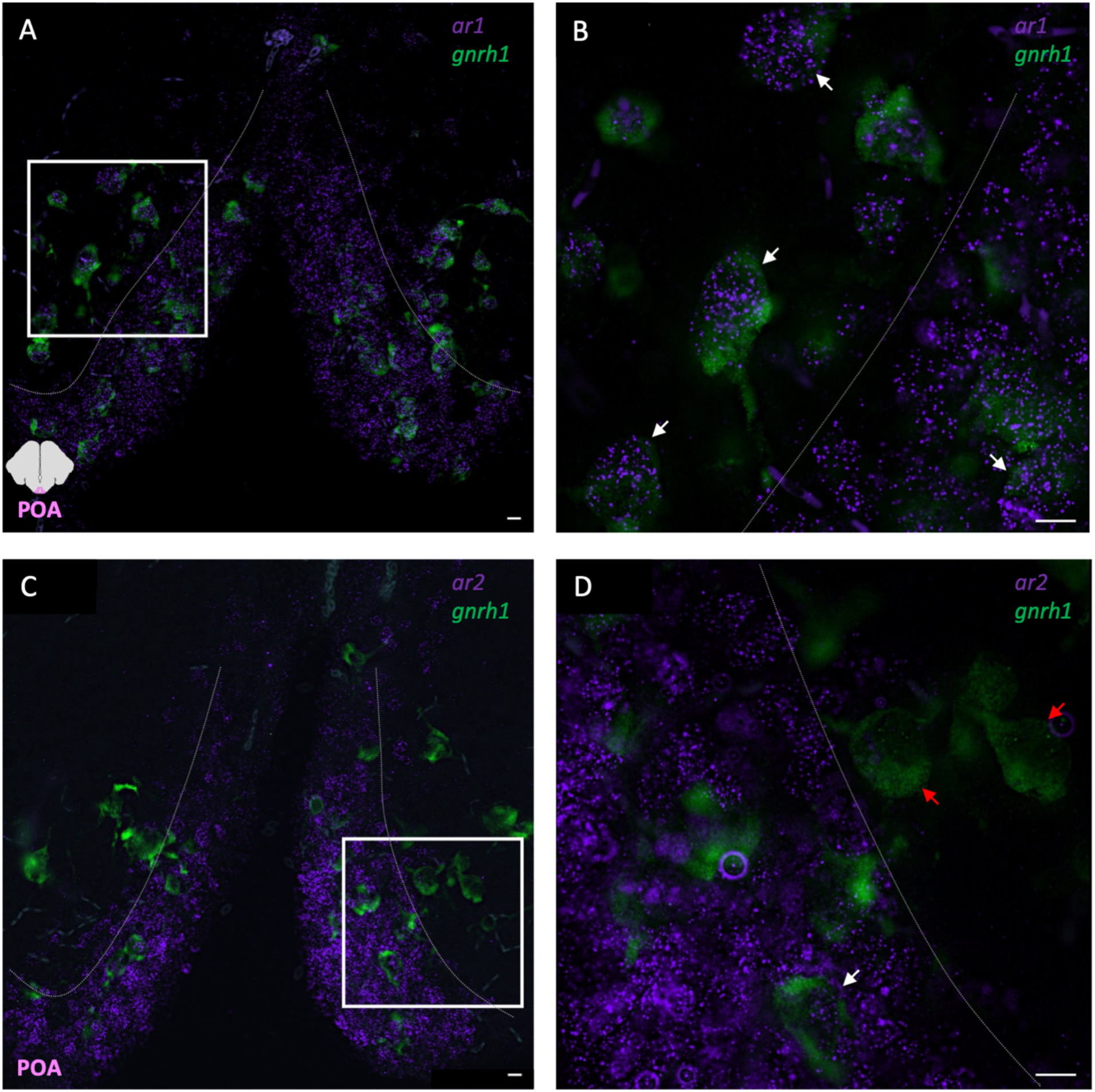
Expression pattern of *ar1* and *ar2* genes in GnRH1 neurons, highlighted using HCR. GnRH1 neurons were labeled using HCR probes (GFP), *ar1* and *ar2* mRNA were localized using HCR probes as well (*ar1* mRNA: panels A, B; *ar2* mRNA: C, D. Cy5). *ar1* gene was expressed in all GnRH1 neurons (A and B, white arrows). *ar2* gene was not expressed in the majority of GnRH1 neurons (D, red arrow) even though a few GnRH1 neurons seem to express this gene (D, white arrow). Dashed lines delineate the AR-rich area of the nPPa. Scale bars, 10 µm.

In order to visualize all three targets at the same time, we adapted a novel protocol consisting of the simultaneous use of HCR probes and antibodies. Thus, we were able to localize both *ar1* and *ar2* transcripts and GnRH1 neurons on the same sections (Fig 4). Again, we confirm that GnRH1 neurons outside of the AR-rich area (delimited by the dashed line, Fig 4) express *ar1* mRNA but not *ar2* mRNA (Fig 4, white and red arrows, respectively). μm.

**Fig 4:**
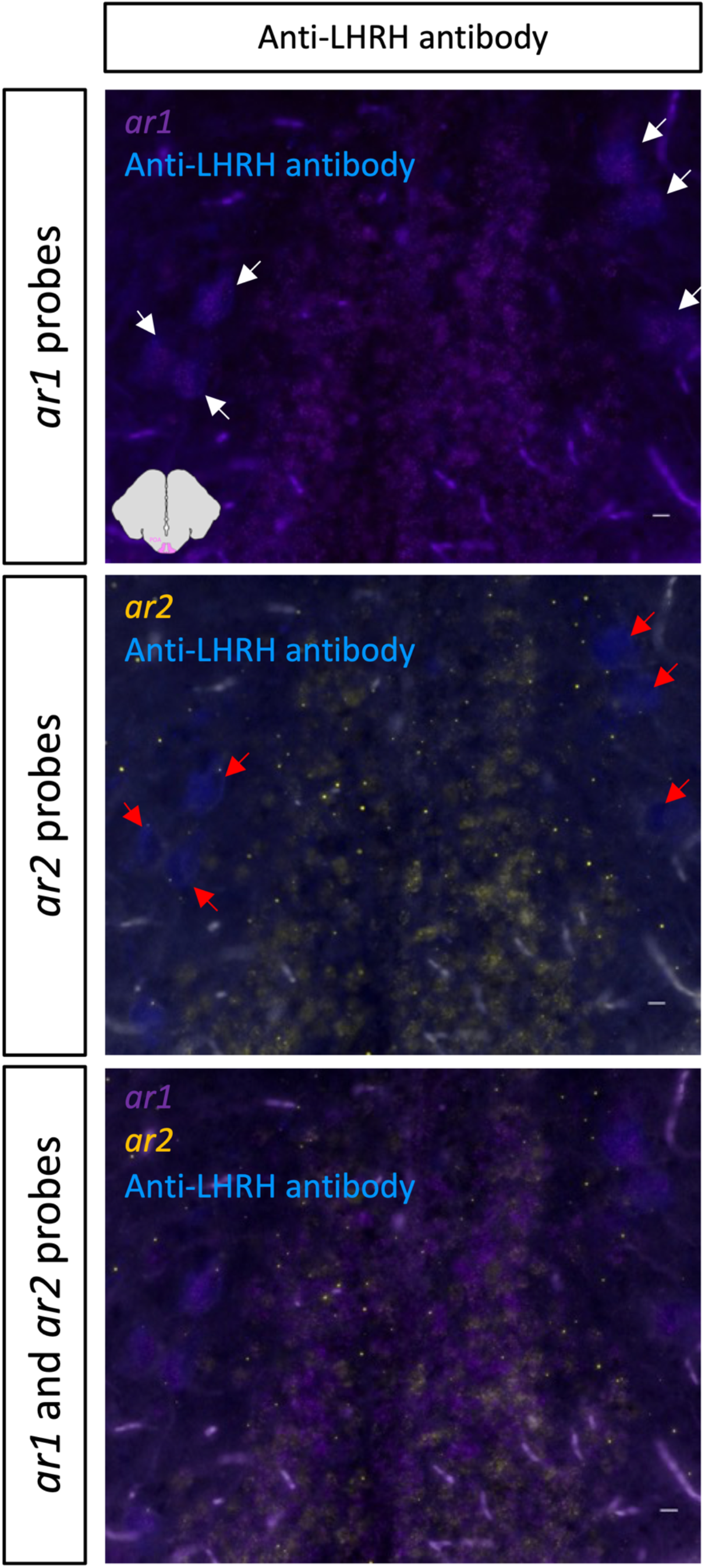
Expression pattern of *ar1* and *ar2* genes in GnRH1 neurons, using a novel protocol combining the use of HCR probes and one antibody. GnRH1 neurons were immunostained using the anti-LHRH antibody (Immunostar, visualized in DAPI), both *ar1* and *ar2* mRNA were localized using the corresponding HCR probes (visualized in Cy5 and YFP, respectively). Pictures show that *ar1* but not *ar2* genes is expressed in GnRH1 neurons (white and red arrows, respectively). Scale bars, 10 μm.

We then investigated the presence of *ar1* and *ar2* transcripts in GnRH2 and GnRH3 neurons, using the two alternate slide series that were processed using HCR probes. In GnRH2 neurons localized in the nMLF, *ar1* and *ar2* transcripts could be seen in only a very few instances (see white arrows in Fig 5 A, C) in both male and female brains. In GnRH3 neurons localized in the olfactory bulb, *ar1* and *ar2* transcripts were both found to be expressed (see white arrows in Fig 5 B, D) in both sexes.

**Fig 5:**
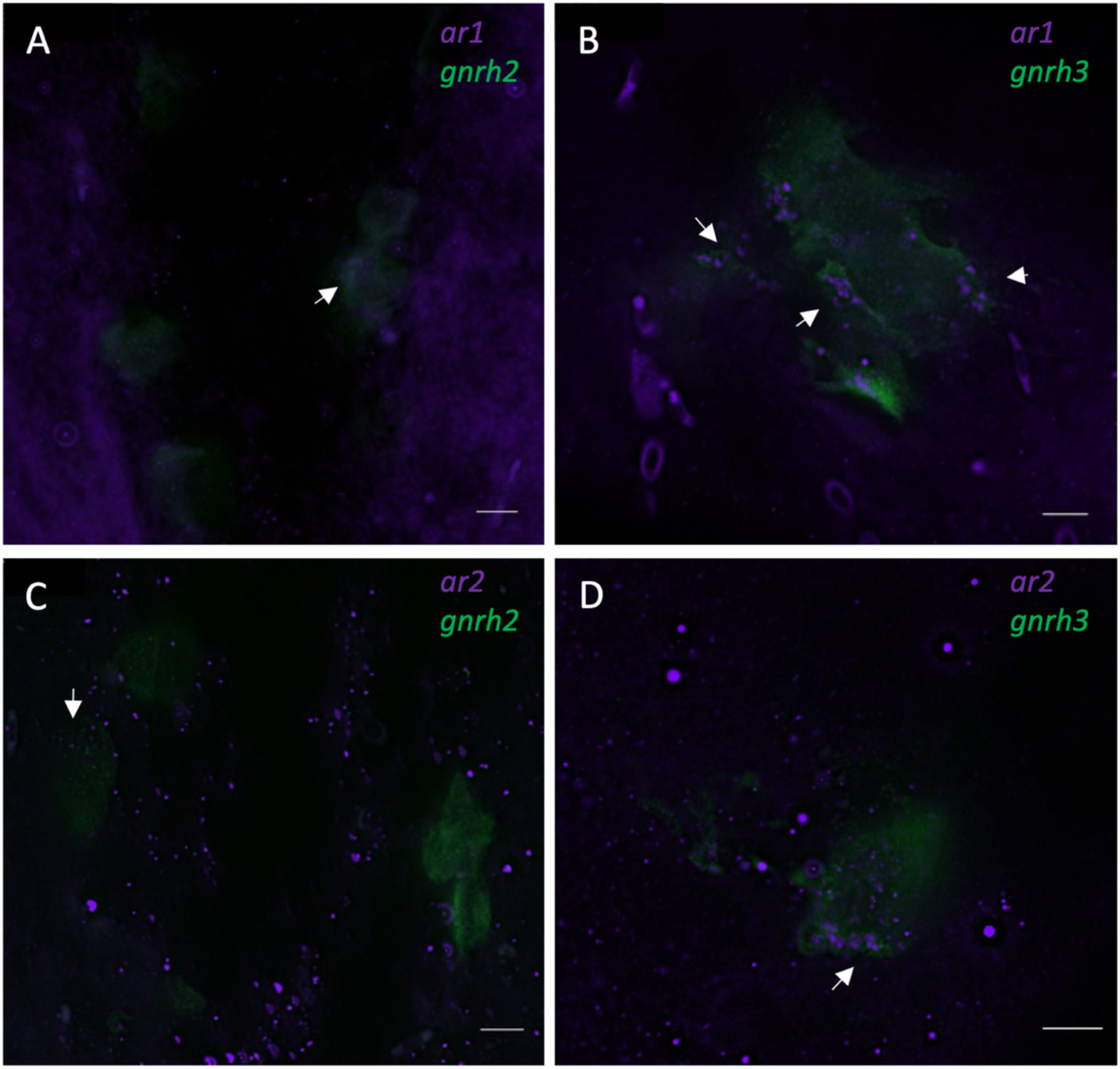
Expression of *ar1* and *ar2* genes (top and bottom panels, respectively) in GnRH2 and GnRH3 neurons (left and right panels, respectively). All transcripts were identified using HCR probes (*ar1* and *ar2* in Cy5; *gnrh2* and *gnrh3* in GFP). Scale bars, 10 μm.

## 4 Discussion

In the present study, we sought to determine whether GnRH neurons express *ar1* and *ar2* genes in the brain of *A. burtoni*. For this purpose, we wished to investigate both the presence of *ar1* and *ar2* transcripts and AR proteins in all three GnRH neuron subpopulations. We show that in GnRH1 neurons, *ar1* and *ar2* genes show a different expression pattern. Indeed, while all GnRH1 neurons expressed the *ar1* gene, only a few of them expressed the *ar2* gene, provided they were localized in the AR-rich area of the POA along the third ventricle. GnRH2 neurons were shown to express *ar1* and *ar2* genes very scarcely. GnRH3 neurons were found to express both *ar1* and *ar2* genes. In a desire to investigate the presence of ARα and ARβ receptor proteins in GnRH neurons, we used custom anti-ARα and anti-ARβ antibodies. Unfortunately, these custom antibodies did not pass the validation process, meaning they were not specifically recognizing their target proteins. Nevertheless, we thoroughly validated an anti-LHRH antibody for use in our species that localizes all three forms of GnRH peptides.

Antibodies directed against AR have been successfully used for many years in mammals 40 which possess only one *ar* gene encoding a single AR protein. In most teleosts however, the whole-genome duplication gave rise to two *ar* gene paralogs (*ar1* and *ar2*), which encode two different proteins (ARα and ARβ) ^31–33^ Both receptors share more than 58% of identity, which complicates the generation of antibodies that specifically recognizes both isoforms. In the present study, we investigated the specificity both anti-ARα and anti-ARβ custom antibodies using two complementary approaches: a genetic strategy, where antibody signal was detected in AR knockouts, and western blotting ^41,42^ Unfortunately, both approaches revealed that these custom antibodies recognize off-target proteins and thus, do not specifically recognize ARα and ARβ proteins. Other attempts to specifically localize both ARs in teleosts using antibodies have been unsuccessful ^43^. This had led some to use antibodies specific to the human AR, an approach that also lacks specificity in regards to novel teleost ARs as well as the presence of off-target signal ^44^.

Since obtaining specific anti-AR antibodies in teleosts is difficult, the generation of a reporter line for both ARα and ARβ might be helpful. In medaka for example, two knock-in lines have been generated to help localize both ARα and ARβ isoforms. Under the control of each *ar* promoter, both an AR-FLAG and a P2A-mClover3 are produced, which can in turn be localized using an anti-FLAG antibody and an anti-GFP antibody, respectively ^45^ Similarly in mice, Shah et al. (2004) 46 modified the *ar* gene such that cells expressing *ar* co-expressed two other molecules: the nuclear targeted lacZ (nLacZ) and placental alkaline phosphatase (PLAP). Thanks to this genetic construction, both nuclei and neuronal processes of *ar*-expressing cells can be labelled, making it possible to investigate neuronal projections of these cells. With the establishment of successful and efficient genome editing technologies available in *A. burtoni*, transgenic approaches for detecting distinct AR proteins are warranted.

We thoroughly validated an anti-GnRH antibody that recognizes all three GnRH forms in *A. burtoni*. To validate this antibody, we first compared the immunostaining to that obtained with an anti-eGFP antibody in our GnRH1:eGFP transgenic strain ^36^ Signals were identical, confirming the anti-LHRH antibody effectively stained GnRH1 neurons. To further confirm this anti-LHRH antibody also specifically recognized GnRH2 and GnRH3 peptides, we pre-incubated it with the target proteins, which resulted in the disappearance of immunolabelling. We therefore conclude that this antibody specifically labels all GnRH forms in *A. burtoni*, which benefits future investigations into all three GnRH neuron subtypes in this species.

The present study demonstrates the presence of *ar1* transcripts in all GnRH1 neurons in both male and female *A. burtoni*. Interestingly, *ar2* transcripts could be observed in just a few GnRH1 neurons, specifically ones located in the AR-rich area of the nPPA along the ventricle. These findings suggest there may exist two types of genetically-defined GnRH1 neurons: ones that only express *ar2* and ones that express both *ar1* and *ar2*. Additionally, these genetically-defined GnRH1 neurons appear to be topographical because *ar2* expression in GnRH1 neurons is restricted to the nPPA along the ventricle. Future work investigating whether *ar1+* and *ar1/2+* GnRH1 neurons functionally differ may yield important insights into the control of reproductive physiology and plasticity by GnRH1 in cichlids and other teleost fishes.

The results from Harbott and colleagues on expression of both *ar* genes in GnRH1 neurons, however, differ 34. This discrepancy in our results might be because in their study, Harbott et al. did not image the brain sections using z-stacks; therefore it might be challenging to judge whether or not signals from both *gnrh* and *ar* transcripts belonged to the same plane. In another study, Ogawa et al. (2020) used qPCR to show that all GnRH neurons subtypes expressed *ar2* but not *ar1* gene in Nile tilapia ^19.^ These findings also contrast with ours, suggesting there may be robust species differences in the expression of *ar1* and *ar2* within GnRH neurons between *A. burtoni* and Nile tilapia. To disentangle the cause of these differences across studies requires the use of standardized approaches for tissue collection and histology and normalizing reproductive state and sex of subjects used.

The detection of *ar* genes within GnRH neurons in *A. burtoni* suggests the direct modulation of GnRH neurons by androgens in this species is possible. However, Kitahashi et al. (2005) cloned the promoters of all GnRH genes in Nile tilapia and, using bioinformatics analysis, showed that none of *gnrh1, gnrh*2 and *gnrh3* gene promoters contained putative binding sites for androgen receptors. Additionally, they showed that even though *gnrh1* and *gnrh2* genes possess putative estrogen binding sites, the *gnrh3* gene seem devoid of it 47 These findings led to the authors to hypothesize that effects of androgen on these GnRH neuronal populations would be exerted via 1) aromatization of testosterone and thus, action on estrogen receptors or 2) the action of upstream androgen-sensitive neurons as it is the case in mice 48. Indeed, some studies showed that E2 treatments could stimulate the HPG axis in fish. For example, administration of E2 to castrated male tilapia increased the *gnrh1* mRNA level compared to control ^49^ Similarly, administration of E2 in both female Nile tilapia and male black porgy (*Acanthopagrus Schegeli*) resulted in the increased mRNA level of GnRH receptor 1 (*gnrh-r1*) *in vivo* and *in vitro* ^50–52^

One explanation, however, is that androgens affect GnRH neurons through non-genomic pathways. The classical signaling pathway of steroids (i.e the genomic pathway) takes place inside the nucleus and is transcription-dependent, which involves protein synthesis which usually occurs within a few hours. On the contrary, the non-classical (i.e the non-genomic) signaling pathways take place outside of the nucleus and do not necessarily involve gene expression.

Typically, in the non-classical mechanism, steroids bind to membrane receptors that activates intracellular pathways in a matter of minutes ^53,54^. Interestingly, in *A. burtoni*, as soon as 20 minutes after the opportunity to rise in social rank, males show both an increase in testosterone and 11-KT and an up-regulation of *egr-1* expression in GnRH1 neurons ^10^. This activation of GnRH1 neurons finally ends in an increase of *gnrh1* mRNA levels, stimulating the whole HPG axis in ascending males ^55^. Thus, one could hypothesize that non-genomic effects of androgens could be exerted via ARα and/or ARβ in GnRH1 neurons, ultimately leading to the activation of the whole HPG axis. Links between androgens and *egr-1* expression has not been investigated in the other GnRH neuron subtypes in *A. burtoni*, but the hypothesis that androgens control all three GnRH neurons is testable.

We also showed that GnRH3 neurons express both *ar1* and *ar2* genes. This finding contradicts the results of Harbott et al. (2007) who did not localize any *ar* transcripts in the olfactory bulbs of *A. burtoni* 34. A study on castrated Nile tilapia males showed that testosterone injection resulted in the increased number of *gnrh3* transcripts 17 which could not be explained by testosterone aromatization since the *gnrh3* gene promoter do not possess AR nor ER binding sites 47. This idea was further confirmed by two important studies showing that in adult tilapia females, injection with the non-aromatizable androgen 11-KT significantly increased the number of *gnrh3* transcripts and stimulated neurogenesis of GnRH3 neurons. Moreover, both studies showed that injection with E2 did not have any of these effects on GnRH3 neurons ^27, 28^.

Therefore, our findings indicate that androgens could act directly on GnRH3 neurons in the olfactory bulb although actions via another androgen-responsive neuronal population, or via a non-genomic action in GnRH3 neurons themselves, cannot be ruled out ^54,56,57^. The differences between the current study and that of Harbott et al., therefore, may be due to sensitivity in signal detection, where fluorescence based HCR might be better at detecting low expressing genes.

## Conclusions

Our findings indicate that all GnRH subpopulations express both *ar1* and *ar2* to different extents. More precisely, the hypophysiotropic GnRH1 neuron population which regulates the HPG axis, strongly express the *ar1* gene. Interestingly, just a few of these GnRH1 neurons express the *ar2* gene. We postulate from this observation that there are distinct, genetically-defined GnRH1 neuron subtypes in the *A. burtoni* brain. Both *ar* genes were seen to be expressed in GnRH2 neurons in very few instances. Lastly, both *ar* genes were expressed in GnRH3 neurons. The mode of action of androgen on these neurons remains unknown, since all three GnRH subtypes apparently lack AR binding sites, but we hypothesize the regulatory mechanism may involve non-genomic signaling.. Overall, these results provide a foundation for future studies to disentangle the androgenic control of GnRH neuron plasticity and reproductive physiology in *A. burtoni* and other teleosts.

## Author contributions

Mélanie Dussenne: conceptualization; investigation; data curation; validation; visualization; writing - original draft; writing - review and editing.

Beau A. Alward: conceptualization; funding acquisition; resources; supervision; writing - review and editing.

## Acknowledgements

We thank Dr Kathleen Munley, Mariana S. Lopez and Megan Howard for the optimization of the HCR protocol. We are very grateful to Dr Colin Haile and Carlos Lopez Arteaga for their precious help regarding the western blotting. We thank Megan Lean for preliminary discussions on the experiment.

## Data availability statement

All of the findings supporting the claims made in this manuscript are available on the figures in the main text and present in the Supplementary Materials file.

## References

1. Sapolsky RM. The Influence of Social Hierarchy on Primate Health. Science. 2005;308:648– 652.

2. Fernald RD. Social Control of the Brain. Annu Rev Neurosci. 2012;35:133–151.

3. Fernald RD, Maruska KP. Social information changes the brain. Proc Natl Acad Sci. 2012;109:17194–17199.

4. Maruska KP. Social Transitions Cause Rapid Behavioral and Neuroendocrine Changes. Integr Comp Biol. 2015;55:294–306.

5. Maruska KP. Social regulation of reproduction in male cichlid fishes. Gen Comp Endocrinol. 2014;207:2–12.

6. Maruska KP, Anselmo CM, King T, et al. Endocrine and neuroendocrine regulation of social status in cichlid fishes. Horm Behav. 2022;139:105110.

7. Munley KM, Alward BA. Control of social status by sex steroids: insights from teleost fishes. Mol Psychol Brain Behav Soc. 2023;2:21.

8. Parikh V, Clement T, Fernald R. Androgen level and male social status in the African cichlid, Astatotilapia burtoni. Behav Brain Res. 2006;166:291–295.

9. Maruska KP, Fernald RD. Behavioral and physiological plasticity: Rapid changes during social ascent in an African cichlid fish. Horm Behav. 2010;58:230–240.

10. Burmeister SS, Jarvis ED, Fernald RD. Rapid behavioral and genomic responses to social opportunity. PLoS Biol. 2005;3:1996–2004.

11. Fernald RD. Cognitive skills needed for social hierarchies. Cold Spring Harb Symp Quant Biol. 2014;79:229–236.

12. Desjardins JK, Hofmann HA, Fernald RD. Social context influences aggressive and courtship behavior in a cichlid fish. PLoS ONE;7. Epub ahead of print 2012. DOI: 10.1371/journal.pone.0032781.

13. Zohar Y, Zmora N, Trudeau VL, et al. A half century of fish gonadotropin-releasing hormones: Breaking paradigms. J Neuroendocrinol. 2022;34:e13069.

14. White SA, Nguyen T, Fernald RD. Social regulation of gonadotropin-releasing hormone. J Exp Biol. 2002;205:2567–2582.

15. Greenwood AK, Fernald RD. Social Regulation of the Electrical Properties of Gonadotropin-Releasing Hormone Neurons in a Cichlid Fish (Astatotilapia burtoni)1. Biol Reprod. 2004;71:909–918.

16. White SA, Kasten TL, Bond CT, et al. Three gonadotropin-releasing hormone genes in one organism suggest novel roles for an ancient peptide. Proc Natl Acad Sci U S A. 1995;92:8363–8367.

17. Soga T, Sakuma Y, Parhar IS. Testosterone differentially regulates expression of GnRH messenger RNAs in the terminal nerve, preoptic and midbrain of male tilapia. Mol Brain Res. 1998;60:13–20.

18. Parhar IS, Soga T, Sakuma Y. Thyroid hormone and estrogen regulate brain region-specific messenger ribonucleic acids encoding three gonadotropin-releasing hormone genes in sexually immature male fish, Oreochromis niloticus. Endocrinology. 2000;141:1618–1626.

19. Ogawa S, Parhar IS. Single-cell gene profiling reveals social status-dependent modulation of nuclear hormone receptors in gnrh neurons in a male cichlid fish. Int J Mol Sci;21. Epub ahead of print 2020. DOI: 10.3390/ijms21082724.

20. Zohar Y, Muñoz-Cueto JA, Elizur A, et al. Neuroendocrinology of reproduction in teleost fish. Gen Comp Endocrinol. 2010;165:438–455.

21. Uchida H, Ogawa S, Harada M, et al. The olfactory organ modulates gonadotropin-releasing hormone types and nest-building behavior in the tilapia Oreochromis niloticus. J Neurobiol. 2005;65:1–11.

22. Ogawa S, Akiyama G, Kato S, et al. Immunoneutralization of gonadotropin-releasing hormone type-III suppresses male reproductive behavior of cichlids. Neurosci Lett. 2006;403:201–205.

23. Okuyama T, Yokoi S, Abe H, et al. A Neural Mechanism Underlying Mating Preferences for Familiar Individuals in Medaka Fish. Science. 2014;343:91–94.

24. Umatani C, Oka Y. Multiple functions of non-hypophysiotropic gonadotropin releasing hormone neurons in vertebrates. Zool Lett. 2019;5:1–10.

25. Francis RC, Jacobson B, Wingfield JC, et al. Castration Lowers Aggression but not Social Dominance in Male Haplochromis burtoni (Cichlidae). Ethology. 1992;90:247–255.

26. Soma KK, Francis RC, Wingfield JC, et al. Androgen Regulation of Hypothalamic Neurons Containing Gonadotropin-Releasing Hormone in a Cichlid Fish: Integration with Social Cues. Horm Behav. 1996;30:216–226.

27. Narita Y, Tsutiya A, Nakano Y, et al. Androgen induced cellular proliferation, neurogenesis, and generation of GnRH3 neurons in the brain of mature female Mozambique tilapia. Sci Rep. 2018;8:16855.

28. Kuramochi A, Tsutiya A, Kaneko T, et al. Sexual dimorphism of gonadotropin-releasing hormone type-III (GnRH3) neurons and hormonal sex reversal of male reproductive behavior in Mozambique tilapia. Zoolog Sci. 2011;28:733–9.

29. Brunet FG, Crollius HR, Paris M, et al. Gene Loss and Evolutionary Rates Following Whole-Genome Duplication in Teleost Fishes. Mol Biol Evol. 2006;23:1808–1816.

30. Glasauer SMK, Neuhauss SCF. Whole-genome duplication in teleost fishes and its evolutionary consequences. Mol Genet Genomics. 2014;289:1045–1060.

31. Ogino Y, Kuraku S, Ishibashi H, et al. Neofunctionalization of Androgen Receptor by Gain-of-Function Mutations in Teleost Fish Lineage. Mol Biol Evol. 2016;33:228–244.

32. Munley KM, Hoadley AP, Alward BA. A phylogenetics-based nomenclature system for steroid receptors in teleost fishes. Gen Comp Endocrinol. 2024;347:114436.

33. Douard V, Brunet F, Boussau B, et al. The fate of the duplicated androgen receptor in fishes: a late neofunctionalization event? BMC Evol Biol. 2008;8:336.

34. Harbott LK, Burmeister SS, White RB, et al. Androgen receptors in a cichlid fish, Astatotilapia burtoni: Structure, localization, and expression levels. J Comp Neurol. 2007;504:57–73.

35. Fernald RD, Hirata NR. Field study of Haplochromis burtoni: habitats and co-habitant. Environ Biol Fishes. 1977;2:299–308.

36. Ma Y, Juntti SA, Hu CK, et al. Electrical synapses connect a network of gonadotropin releasing hormone neurons in a cichlid fish. Proc Natl Acad Sci. 2015;2:201421851.

37. Alward BA, Laud VA, Skalnik CJ, et al. Modular genetic control of social status in a cichlid fish. Proc Natl Acad Sci U S A. 2020;117:28167–28174.

38. Dussenne M, Gennotte V, Rougeot C, et al. Hormones and Behavior Consequences of temperature-induced sex reversal on hormones and brain in Nile tilapia (Oreochromis niloticus). Horm Behav. 2020;121:104728.

39. Ćorić A, Stockinger AW, Schaffer P, et al. A Fast And Versatile Method for Simultaneous HCR, Immunohistochemistry And Edu Labeling (SHInE). Integr Comp Biol. 2023;63:372– 381.

40. Pomerantz MM, Li F, Takeda DY, et al. The androgen receptor cistrome is extensively reprogrammed in human prostate tumorigenesis. Nat Genet. 2015;47:1346–1351.

41. Uhlen M, Bandrowski A, Carr S, et al. A proposal for validation of antibodies. Nat Methods. 2016;13:823–827.

42. Pillai-Kastoori L, Heaton S, Shiflett SD, et al. Antibody validation for Western blot: By the user, for the user. J Biol Chem. 2020;295:926–939.

43. Takeo J, Yamashita S. Immunohistochemical localization of rainbow trout androgen receptors in the testis. Fish Sci. 2001;67:518–523.

44. Gelinas D, Callard GV. Immunolocalization of Aromatase- and Androgen Receptor-Positive Neurons in the Goldfish Brain. Gen Comp Endocrinol. 1997;106:155–168.

45. Ogino Y, Ansai S, Watanabe E, et al. Evolutionary differentiation of androgen receptor is responsible for sexual characteristic development in a teleost fish. Nat Commun. 2023;14:1428.

46. Shah NM, Pisapia DJ, Maniatis S, et al. Visualizing Sexual Dimorphism in the Brain. Neuron. 2004;43:313–319.

47. Kitahashi T, Sato H, Sakuma Y, et al. Cloning and functional analysis of promoters of three GnRH genes in a cichlid. Biochem Biophys Res Commun. 2005;336:536–543.

48. Zwain IH, Arroyo A, Amato P, et al. A Role for Hypothalamic Astrocytes in Dehydroepiandrosterone and Estradiol Regulation of Gonadotropin-Releasing Hormone (GnRH) Release by GnRH Neurons. Neuroendocrinology. 2002;75:375–383.

49. Parhar IS, Soga T, Sakuma Y. Thyroid Hormone and Estrogen Regulate Brain Region-Specific Messenger Ribonucleic Acids Encoding Three Gonadotropin-Releasing Hormone Genes in Sexually. Endocrinology. 2000;141:1618–1626.

50. Levavi-Sivan B, Biran J, Fireman E. Sex Steroids Are Involved in the Regulation of Gonadotropin-Releasing Hormone and Dopamine D2 Receptors in Female Tilapia Pituitary1. Biol Reprod. 2006;75:642–650.

51. Lin C-J, Wu G-C, Lee M-F, et al. Regulation of two forms of gonadotropin-releasing hormone receptor gene expression in the protandrous black porgy fish, Acanthopagrus schlegeli. Mol Cell Endocrinol. 2010;323:137–146.

52. Lin CJ, Wu GC, Dufour S, et al. Activation of the brain-pituitary-gonadotropic axis in the black porgy Acanthopagrus schlegelii during gonadal differentiation and testis development and effect of estradiol treatment. Gen Comp Endocrinol. 2019;281:17–29.

53. Thomas P. Membrane Androgen Receptors Unrelated to Nuclear Steroid Receptors. Endocrinology. 2019;160:772–781.

54. Hammes SR, Davis PJ. Overlapping nongenomic and genomic actions of thyroid hormone and steroids. Best Pract Res Clin Endocrinol Metab. 2015;29:581–593.

55. Maruska KP, Fernald RD. Social Regulation of Male Reproductive Plasticity in an African Cichlid Fish. 2013;53:938–950.

56. Thomas P. Rapid steroid hormone actions initiated at the cell surface and the receptors that mediate them with an emphasis on recent progress in fish models. Gen Comp Endocrinol. 2012;175:367–383.

57. Foradori CD, Weiser MJ, Handa RJ. Non-genomic actions of androgens. Front Neuroendocrinol. 2008;29:169–181.

