## Supplementary Material for "Expression of novel androgen receptors in three GnRH neuron subtypes in the cichlid brain"

**Affiliations**
a *University of Houston, Department of Psychology, United States of America*
b *University of Houston, Department of Biology and Biochemistry, United States of America*

**ORCID ID:**
Mélanie Dussenne: 0000-0002-1903-883X
Beau A. Alward: 0000-0002-1403-5277

### Supplementary Information

Methods and Results

**1. Western blotting**
Several tissues were used for western blotting: testes and brain of a WT male; ovaries and testes of Bd5 KO individuals; ovaries and liver of Ad50 KO females; testes of a Ad50 KO male. These tissues were selected because they were shown to express both ar1 and ar2 genes in WT fish [1,2].
These tissues of interest were dissected on ice and weighted. Ice cold lysis buffer (RIPA buffer, 89901, Thermo, containing Halt Protease and Phosphatase Inhibitor Cocktail 100x, 78440, Thermo) was added to each sample (1 ml of lysis buffer for 50 mg of tissue). Samples were homogenized on ice, centrifuged at 10,000 g for 5 min at 4°C, supernatants were collected in a clean tube and stored at -20°C until further use.
For each sample, protein concentration was measured using a modified Lowry assay according to the manufacturer's instructions (*RC DC*™ Protein Assay Kit I #5000121, Biorad). Then, 20 µg of protein was sampled from each tube and added to an equal volume of 2x Laemmli buffer (1610737, Biorad). Samples were boiled at 95°C for 5 min and centrifuged at 16,000 g for 1 min at 4°C. Protein were then separated using a one-dimensional SDS-polyacrylamide gel electrophoresis using 7.5% acrylamide gel (4561023, Biorad). Proteins were transferred onto a nitrocellulose membrane (iBlot gel transfer device, IB21001; iBlot transfer stacks, 23002, Thermo), which was subsequently rinsed in pure water and stained with Ponceau S staining solution (A40000279, Thermo) to check for transfer quality. The Ponceau S stain was then rinsed off with TBST washes (3 x 5 min) and the membrane was blocked for 5 min (EveryBlot Blocking Buffer, 12010020, Biorad). The blot was then incubated in the primary antibody solution for 1 hour at RT (anti-ARα custom antibody, Aves, 1/1000 in everyblot blocking buffer,

Biorad), rinsed with TBST (4 x 5 min, RT) and incubated in the secondary antibody solution for 1 hour at RT (Goat anti-chicken antibody, HRP conjugated, PA1-28798, Thermo). After additional TBST rinses (4 x 5min, RT), the chemoluminescent substrate (Clarity western substrate, 1705060, Biorad) was applied to the blot for 5 min. The blot was imaged (C-DiGit blot scanner, Li-Cor) after which it was rinsed (TBST, 4 x 5 min) and stripped (30 min, RT, with Restore Plus Stripping Buffer, 46430, Thermo). After additional TBST rinses, the blot was re-developped with the anti-ARβ custom antibody (Aves, 1/1000) and the same secondary antibody as previously.


1. Chakraborty SB, Banerjee S, Chatterjee S. 2011 Increased androgen receptor expression in muscle tissue contributing to growth increase in androgen-treated Nile tilapia. *Aquac. Int.* **19**, 1119–1137.
2. Schuppe ER, Pradhan DS, Thonkulpitak K, Drilling C, Black M, Grober MS. 2017 Sex differences in neuromuscular androgen receptor expression and sociosexual behavior in a sex changing fish. *PLoS ONE* **12**.


#### Supplementary tables

Supplementary Table 1: Primer sequences used for the determination of fish genotype

| Strain | Primer name | Direction | Sequence |
|---|---|---|---|
| Ad50 mutant | AR1FlankF3 | Forward | 5′-CCC AGT GCA CTC TAA CTC CG-3′ |
| | AR1FlankR3 | Reverse | 5′-TTT AAG GGT ACG ACC TCG GC-3′ |
| Bd5 mutant | AR2FlankF1 | Forward | 5′-CCA TCC CAC CTC CAA GAG TC-3′ |
| | AR2FlankR3 | Reverse | 5′-GAG GAC AGG CCG ATG ATG AA-3′ |
| gfp:gnrh1 transgenic | GnRH1genoFwd | Forward | 5′- GCA GCA GAC TTC ACA AAG GAC AG-3′ |
| | EGFPgenoRev | Reverse | 5′- AGC TTG CCG TAG GTG GCA TC-3′ |
